## Supplementary Figures and Table for "Patchy Striatonigral Neurons Modulate Locomotor Vigor in Response to Environmental Valence"

### Supplementary Figures, Figure legends, and Table

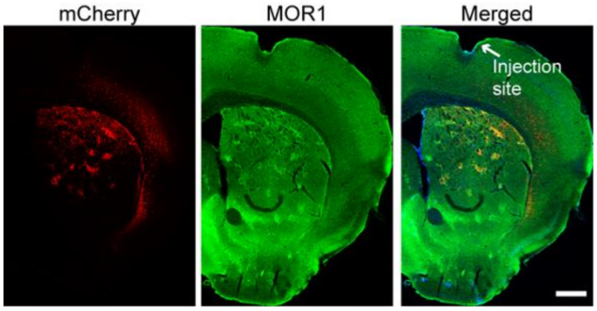

**Fig. S1 Patchy distribution of mCherry-positive cells in the dorsal striatum of *Sepw1-Cre* mice.**

Representative images show mCherry (red) and MOR1 (green) immunostaining in the dorsal striatum of *Sepw1-Cre* mice following stereotactic injection of AAV-EF1a-DIO-mCherry. Scale bar: 500μm.

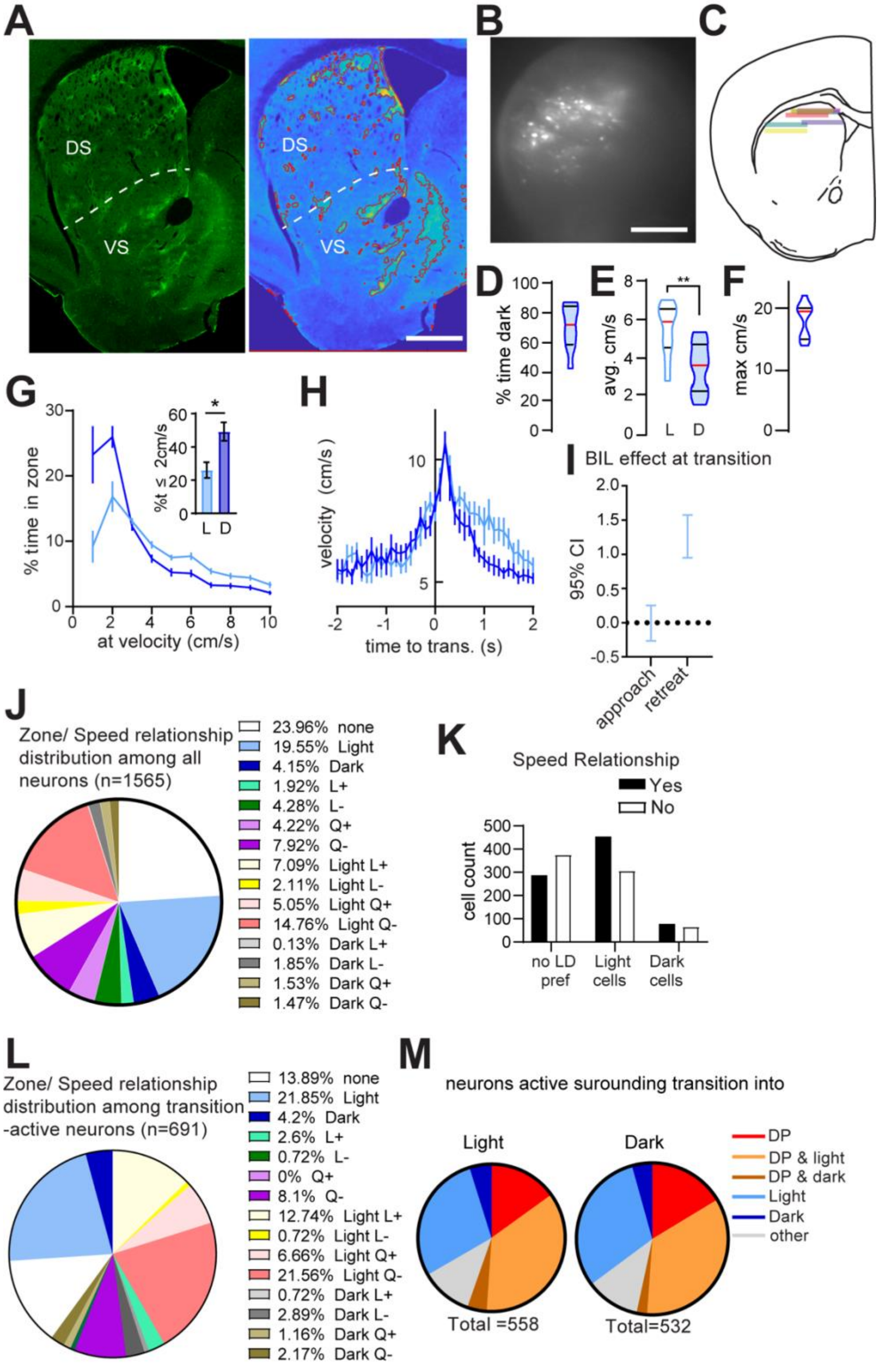

**Fig. S2. MOR1 thresholding, Scope-mounted Light/Dark box behavior and $Ca^{2+}$ imaging supplemental. (A)** Patch ablation was assessed by MOR1-positive area measurements. Example dorsal striatal (DS) and ventral striatal (VS) hemisection stained for MOR1 (left) alongside thresholded image generated in MATLAB (right) used for quantifying MOR1-positive territories. Scale bar: 500μm. **(B)** Example field of view through GRIN lens. Scale bar: 500μm. **(C)** Schematic of lens positions in imaged mice. **(D-L)** N=10 (7M, 3F) scope-mounted mice. **(D)** % time in dark. **(E)** Average speed. Wilcoxon L vs D, **p=0.0098. **(F)** Maximum speed. **(G)** Speed distribution normalized to zone. %time ≤2cm/sec (bar graph) Wilcoxon L vs D, *p=0.0137. **(H)** Transition speed. **(I)** GLME fixed effects 95% CI (cm/s) in brackets. Approach speed is unchanged by BIL [0.2506, -0.26603]. Retreat speed is increased by BIL [1.576, 0.9505]. **(J)** Distribution of zone and/or speed relationships among all recorded neurons. **(K)** Relationship between zone preference and speed encoding $\chi^2$ (3, n=532) =78.63, p<0.0001 **(L)** Distribution of zone and/or speed relationships among all transition-active neurons. **(M)** Distribution of zone and/or deceleration relationships among all transition-active neurons.

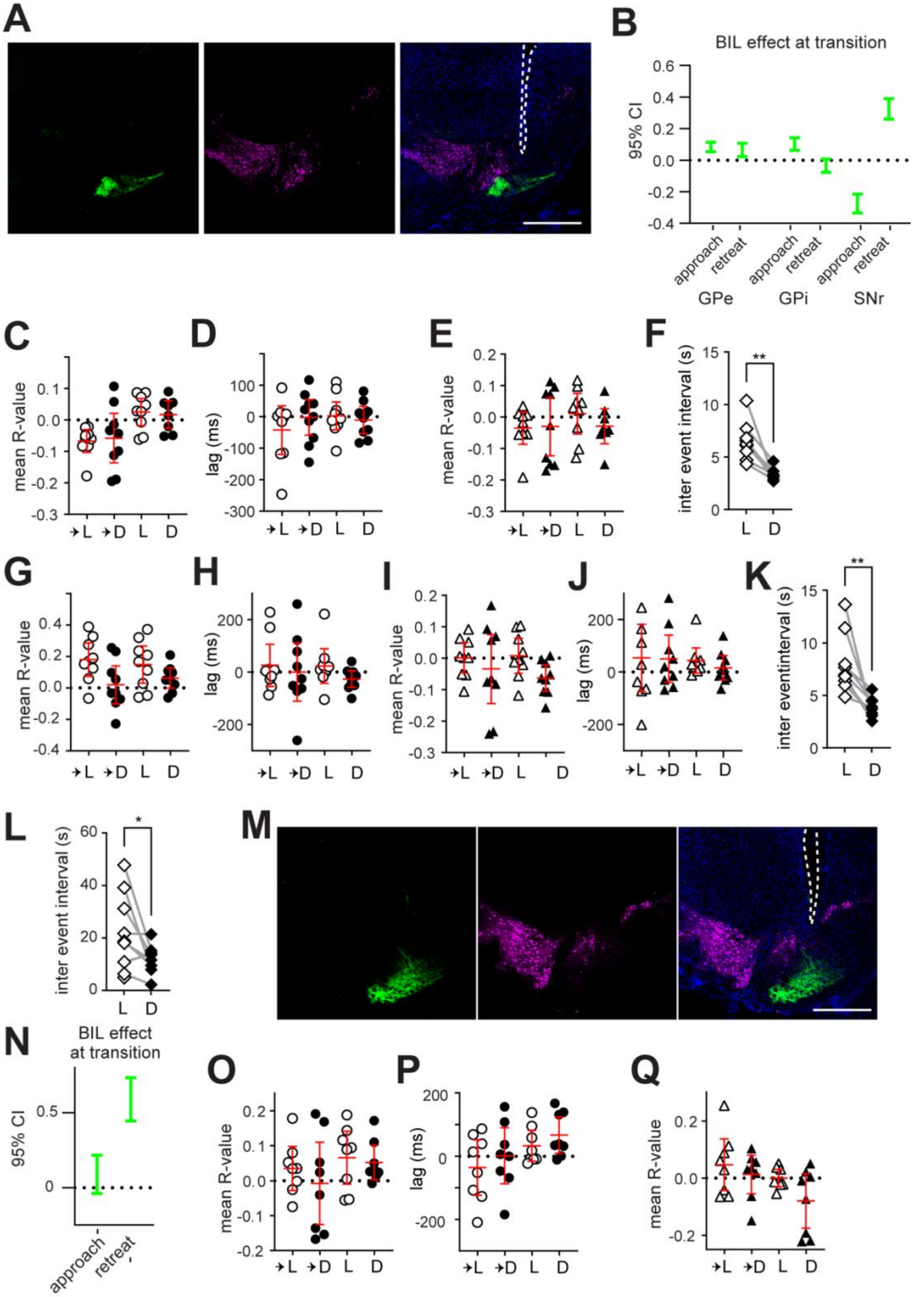
A
B
BIL effect at transition
95% CI
approach
retreat
GPe
GPi
SNr
C
D
E
F
G
H
I
J
K
L
M
N
O
P
Q
mean R-value
lag (ms)
inter event interval (s)
inter eventinterval (s)
→L
→D
L
D
**
*
BIL effect
at transition

**Fig. S3. Fiber photometry supplemental. (A-K)** N=8 (4M, 4F) GPe-implanted, 9 (5M, 4F) GPi-implanted *Sepw1*-Cre mice. **(A)** Representative images show GCaMP8s (green) and TH (magenta) immunostaining. Scale bar: 500μm. **(B)** Transition BIL effect on patchy SPN efferents. GLME fixed effects 95% CI (%ΔF/F) in brackets. **(C-F)** Analysis of patchy SPN efferents in GPe. 95% CI in red. **(C)** For all acceleration events with significant ($p<0.05$) ΔF/F cross-correlation to speed, mean R value per mouse, **(D)** Mean Xcorr lag per mouse. **(E)** For all acceleration events with significant ($p<0.05$) ΔF/F cross-correlation to acceleration, mean R value per mouse. **(F)** Event frequent in either zone, paired t-test, **p=0.0018. **(G-K)** Analysis of patchy SPN efferents in GPi. 95% CI in red. **(G)** For all acceleration events with significant ($p<0.05$) ΔF/F cross-correlation to speed, mean R value per mouse. **(H)** Mean Xcorr lag per mouse. **(I)** For all acceleration events with significant ($p<0.05$) ΔF/F cross-correlation to acceleration, mean R value per mouse. **(J)** Mean Xcorr lag per mouse. **(K)** Event frequent in either zone, paired t-test, **p=0.0035. **(L)** N=9 (5M, 4F) GRIN-implanted *Sepw1*-Cre mice. From net fluorescence collected in dorsal striatum, event frequent in either zone, paired, t-test *p=0.0373. **(M-Q)** N= 8 (4M, 4F) SNr-implanted *Calb1*-Cre mice. **(M)** Representative images show GCaMP8s (green) and TH (magenta) immunostaining. Scale bar: 500μm. **(N)** Transition BIL effect on matrix efferents. GLME fixed effects 95% CI (%ΔF/F) in brackets. **(O-Q)** Analysis of matrix efferents in SNr. 95% CI in red. **(O)** For all acceleration events with significant ($p<0.05$) ΔF/F cross-correlation to speed, mean R value per mouse. **(P)** Mean Xcorr lag per mouse. **(Q)** For all acceleration events with significant ($p<0.05$) ΔF/F cross-correlation to acceleration, mean R value per mouse.

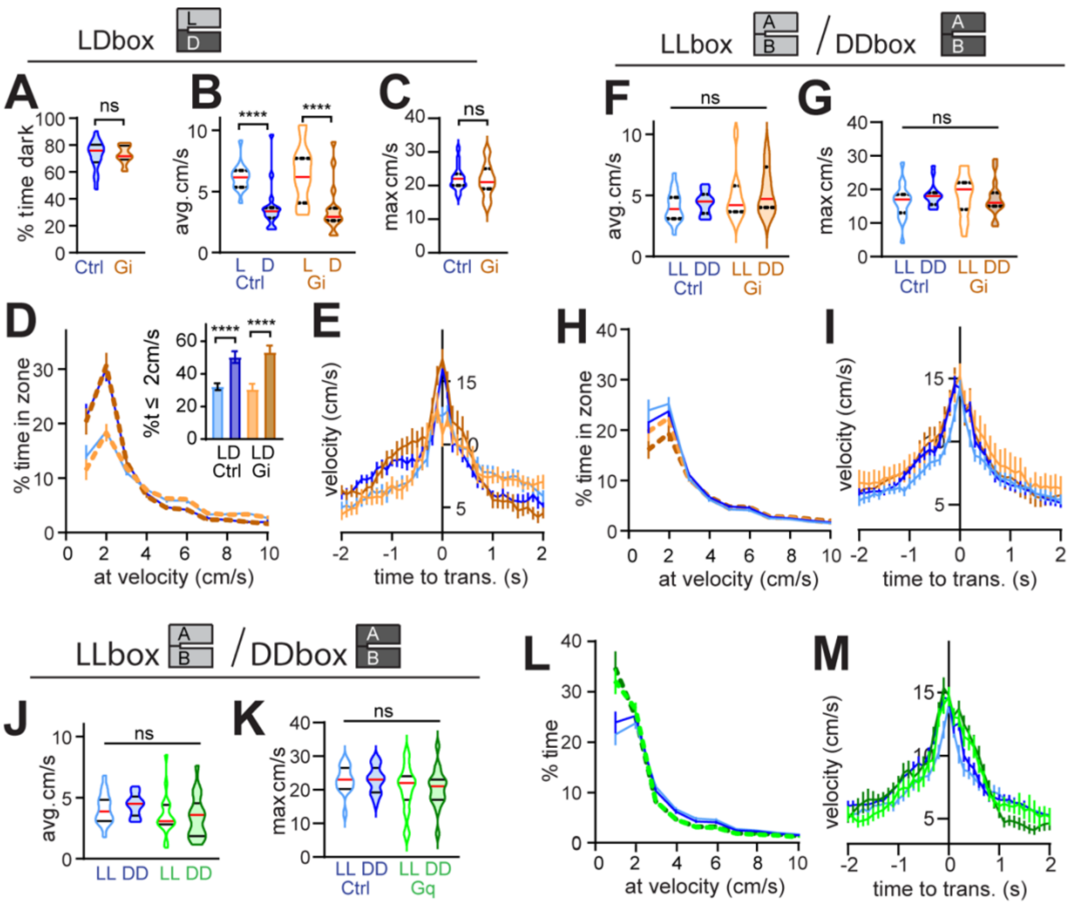

**Fig. S4. Chemogenetics supplemental. (A-E) Gi-mice L/D box**, n=17 (9M, 8F) Ctrl and 15 (8M, 7F) Gi mice. **(A)** % time in the dark. Mann-Whitney, Ctrl vs Gi p=0.8232. **(B)** Average speed. 2way RM ANOVA: group F (1, 30) = 0.03006, p>0.05; light F (1, 30) = 134.3, p<0.0001. **(C)** Maximum speed. 2-tailed Mann Whitney, p=0.715. **(D)** Speed distribution normalized to zone. Insert for % time ≤2cm/sec (bar graph) paired t-test, L vs D Ctrl p<0.0001, Gi p<0.0001, unpaired t-test, Ctrl vs Gi in Light p>0.05, in Dark p>0.05. **(E)** Transition speed. GLME fixed effects 95% CI (cm/s) in brackets. Approach speed is increased by BIL [1.0773, 0.59214] and unchanged by Gi [-1.0473, 1.0196]; Retreat speed is increased by BIL [1.0562, 1.486] and unchanged by Gi [-1.3062, 0.92042]. GLME by group shows approach speed is increased by BIL for control [1.9076, 1.3185] and Gi [2.2463, 1.5553]. **(F-I) Gi-mice LL/DD box**. n= mouse (16 Ctrl, 13 Gi). **(F)** Average speed. 2way RM ANOVA: lighting F (1, 27) = 3651 p=0.0667, group F (1, 27) = 2.994 p>0.05. **(G)** Maximum speed. 2way RM ANOVA: lighting F (1, 27) = 0.05506 p>0.05, group F (1, 27) = 1.402 p>0.05. **(H)** Speed distribution normalized to zone. For %time ≤2cm/sec (bar graph) Wilcoxon LL vs DD Ctrl p>0.05, Gi p=0.0105, Mann-Whitney, Ctrl vs Gi in Light p>0.05, in Dark p>0.05. **(I)** Transition speed. GLME fixed effects 95% CI (cm/s) in brackets. Approach speed is slightly increased by BIA [0.85304, 0.42385], unchanged by Gi [-0.25536, 1.985]; Retreat speed is slightly increased by BIA [0.31052, 0.76904], unchanged by Gi [-0.73445, 1.7233]. **(J-M) Gq-mice LL/DD box**. n= mouse (16 Ctrl, 19 Gq). **(J)** Average speed. 2way RM ANOVA: lighting F (1, 33) = 2.676 p>0.05, group F (1, 33) = 1.343 p>0.05. **(K)** Maximum speed. 2way RM ANOVA: lighting F (1, 33) = 2.683 p>0.05, group F (1, 33) = 0.0008 p>0.05. **(L)** Speed distribution normalized to zone. For %time ≤2cm/sec Wilcoxon LL vs DD Ctrl p>0.05, Gq p>0.05, Mann-Whitney Ctrl vs Gq in LL p=0.0441, in DD p=0.0288. **(M)** Transition speed. GLME fixed effects 95% CI (cm/s) in brackets. Approach speed is increased by BIA [0.85304, 0.42385], unchanged by Gq [-1.1387, 0.89713]; Retreat speed is increased by BIA [0.31052, 0.76904], unchanged by Gq [-1.5338, 0.69961].

**Supplementary Table 1 – Statistical Summary**

| Figure | Test description | Test used | Test statistics | N | N defined as | Mean ± SEM |
|---|---|---|---|---|---|---|
| 1C | DS %MOR1 (Ctrl vs PA) | 2-tailed Mann-Whitney | $p<0.0001$ | n=22 Ctrl; n=30 PA | hemisection. (6-10/region. 3 mice/group) | Ctrl 12.03±0.9643; PA 5.154±0.5749 |
| | VS %MOR1 (Ctrl vs PA) | 2-tailed Mann-Whitney | $p=0.4359$ | n=22 Ctrl; n=30 PA | | Ctrl 21.11±1.357; PA 17.29±1.112 |
| 1E | % time in dark (Ctrl vs PA) | 2-tailed Mann-Whitney | $p=0.3144$ | n=11 Ctrl; n=10 PA | mouse | Ctrl 66.07±3.114; PA 61.6±3.85 |
| 1F | avg. spd cm/s (lighting x group) | 2-way RM ANOVA | $F(1, 19) = 0.06497$, $p=0.8015$ | n=21 (11 Ctrl, 10 PA) | mouse | |
| | avg. spd cm/s (group: Ctrl, PA) | | $F(1, 19) = 5.850$, $p=0.0258$ | | | |
| | avg. spd cm/s (lighting: L, D) | | $F(1, 19) = 22.37$, $p=0.0001$ | | | |
| | avg. spd cm/s (subject) | | $F(19, 19) = 2.849$, $p=0.0138$ | | | |
| | post hoc (Ctrl L-vs-D) | Wilcoxon signed rank | $p=0.001$ | n=11 Ctrl-L; n=11 Ctrl-D | mouse | Ctrl-L 6.01±0.3889; Ctrl-D 3.682±0.3192 |

| | | | | | | |
|---|---|---|---|---|---|---|
| | post hoc (PA L-vs-D) | Wilcox on signed rank | p=0.0371 | n=10 SA-L; n=10 PA-D | mouse | PA-L 8.266±1.322; PA-D 5.673±0.4979 |
| | post hoc (Ctrl-L vs PA-L) | 2-tailed Mann-Whitney | p=0.1971 | n=11 Ctrl; n=10 PA | mouse | see above |
| | post hoc (Ctrl-D vs PA-D) | 2-tailed Mann-Whitney | p=0.0015 | n=11 Ctrl; n=10 PA | mouse | see above |
| 1G | max spd cm/s (Ctrl vs PA) | 2-tailed Mann-Whitney | p=0.0434 | n=11 Ctrl; n=10 PA | mouse | Ctrl 30.36±0.7778; PA 33.3±1.075 |
| 1H | %time ≤2cm/s (Ctrl L-vs-D) | paired ttest | p=0.001 | n=11 Ctrl; n=10 PA | mouse | Ctrl-L 34.72±3; Ctrl-D 52.41±3.441 |
| | %time ≤2cm/s (PA L-vs-D) | paired ttest | p=0.1955 | | | PA-L 29.44±5.285; PA-D 36.54±2.604 |
| | %time ≤2cm/s (Ctrl vs PA in D) | unpaired ttest | p=0.0018 | | | |
| | %time ≤2cm/s (Ctrl vs PA in L) | unpaired ttest | p=0.3847 | | | |
| 1I-J | approach (cm/s) impact of time | GLME of spd uing Matlab | 95% C.I. = [2.3907, 3.1368] | n=630 observations | time points | |
| | approach (cm/s) impact of PA | fitglme'; fixed effects: | 95% C.I. = [0.73218, 3.9241] | | | |

| | | | | | | |
|---|---|---|---|---|---|---|
| | approach (cm/s) impact of BIL | intercept, time, group, BIL; | 95% C.I. = [2.3911, 1.7464] | (11 Ctrl and 10 PA mice at | | |
| | retreat (cm/s) impact of time | random effect: mouse | 95% C.I. = [-1.6015, -0.85885] | at 15 into-D and 15 into-L | | |
| | retreat (cm/s) impact of PA | | 95% C.I. = [2.2344, 2.8761] | time points) | | |
| | retreat (cm/s) impact of BIL | | 95% C.I. = [0.89109, 3.593] | | | |
| | PA approach (cm/s) impact of BIL | GLME of spd uing Matlab | 95% C.I. = [3.347, 2.3376] | n=300 observations | time points | |
| | PA retreat (cm/s) impact of BIL | fitglme'; fixed effects: | 95% C.I. = [3.2809, 4.2788] | (10 mice @15 pnts app/ret) | | |
| | Ctrl approach (cm/s) impact of BIL | intercept, time, BIL; | 95% C.I. = [1.7612, 0.9698] | n=330 observations | time points | |
| | Ctrl retreat (cm/s) impact of BIL | random effect: mouse | 95% C.I. = [1.0784, 1.8056] | (11 mice @15 pnts app/ret) | | |
| 1L | avg. spd cm/s (group: Ctrl, PA) | Mixed-effects analysis | F (1, 12) = 0.1630, p=0.6935 | n=14 (8 Ctrl, 6 PA) | mouse | PA-LL 6.009±0.817; PA-DD 7.026±2.116 |
| | avg. spd cm/s | | F (1, 8) = 0.01655, | | | Ctrl-LL 6.404±1.914; |

| | | | | | | |
|---|---|---|---|---|---|---|
| | (lighting: LL, DD) | | p=0.9008 | | | Ctrl-DD 5.464±1.179 |
| | avg. spd cm/s (group x lighting) | | F (1, 8) = 0.7541, p=0.4105 | | | |
| 1M | max spd cm/s (group: Ctrl, PA) | Mixed-effects analysis | F (1, 11) = 0.05682, p=0.816 | n=13 (7 Ctrl, 6 PA) | mouse | Ctrl-LL 28.86±2.963; Ctrl-DD 25.83±2.428 |
| | max spd cm/s (lighting: LL, DD) | | F (1, 9) = 0.5451, p=0.4791 | | | PA-LL 28.2±2.396; PA-DD 26.83±3.842 |
| | max spd cm/s (group x lighting) | | F (1, 9) = 0.3096, p=0.5915 | | | |
| 1N | %time ≤2cm/s (Ctrl LL-vs-DD) | Wilcoxon signed rank | p=0.8438 | n=7 Ctrl-LL; n=6 Ctrl-DD; | mouse | Ctrl-LL 38.35±7.474; Ctrl-DD 37.09±4.185 |
| | %time ≤2cm/s (PA LL-vs-DD) | Wilcoxon signed rank | p=0.625 | n=5 PA-LL; n=6 PA-DD | | PA-LL 34.54±2.146; PA-DD 39.22±6.015 |
| | %time ≤2cm/s (Ctrl vs PA in DD) | 2-tailed Mann-Whitney | p=0.3939 | | | |
| | %time ≤2cm/s (Ctrl vs PA in LL) | 2-tailed Mann-Whitney | p=0.7551 | | | |
| 1O-P | approach spd (cm/s) impact of time | GLME of spd uing Matlab | 95% C.I. = [2.4259, 3.2878] | n=360 observations | time points | |

| | | | | | | |
|---|---|---|---|---|---|---|
| | approach spd (cm/s) impact of PA | fitglme'; fixed effects: | 95% C.I. = [-2.6119, 4.2595] | (7 LL, 6 DD Ctrl; 5 LL, 6 DD PA | | |
| | approach spd (cm/s) impact of BIA | intercept, time, grp, BIA; | 95% C.I. = [1.0937, -0.3197] | at 15 into-DD, 15 into-LL | | |
| | retreat spd (cm/s) impact of time | random effect: mouse | 95% C.I. = [-1.9249, -1.0011] | time points) | | |
| | retreat spd (cm/s) impact of PA | | 95% C.I. = [-3.708, 3.6125] | | | |
| | retreat spd (cm/s) impact of BIA | | 95% C.I. = [-0.6257, 0.20744] | | | |
| 2E | average ΔF/F in L vs D (Light-neurons) | paired ttest | p<0.0001 | n=760 | neuron | L 0.02657±0.0007633; D 0.1213±0.0004885 |
| | average ΔF/F in L vs D (Dark-neurons) | paired ttest | p<0.0001 | n=143 | neuron | L 0.009896±0.001262; D 0.01948±0.00189 |
| | average ΔF/F in L vs D (Other-neurons) | paired ttest | p=0.016 | n=29 | neuron | L 0.01057±0.000623; D 0.01021±0.0006165 |
| 2I | % light-prefering neurons by mouse | | | n=9 | mouse | 20.95±2.716 |
| | % light&speed neurons by mouse | | | n=9 | mouse | 28.81±3.088 |

| | | | | | | |
|---|---|---|---|---|---|---|
| | % speed neurons by mouse | | | n=9 | mouse | 19.1±2.132 |
| | % dark&speed neurons by mouse | | | n=9 | mouse | 4.655±1.099 |
| | % dark-prefering neurons by mouse | | | n=9 | mouse | 2.412±0.718 |
| 2J | % VL+neurons by mouse | | | n=9 | mouse | 7.81±2.248 |
| | % VL-neurons by mouse | | | n=9 | mouse | 11.20±3.348 |
| | % VQ+ neurons by mouse | | | n=9 | mouse | 9.59±1.703 |
| | % VQ-neurons by mouse | | | n=9 | mouse | 23.95±2.262 |
| | % other types of neurons by mouse | | | n=9 | mouse | 47.44±3.599 |
| 2k | zone/speed encoding relation to speed encoding | Chi Square | X2(3, n=532) =78.63, p<0.0001 | n=532 | neuron | |
| 2M | zone/speed relation to decel. Encoding | Chi Square | X2(1, n=1565) =24.33, p<0.0001 | n=1565 | neuron | |
| & S1N | ΔF/F approach impact of BIL | fitglme'; fixed effects: | 95% C.I. = [0.78166, 0.39086] | | | |

| | | | | | |
|---|---|---|---|---|---|
| | ΔF/F retreat impact of time | intercept, time, BIL; | 95% C.I. = [-0.82892, -0.36778] | (10 mice at 15 into-D and 15 | |
| | ΔF/F retreat impact of BIL | random effect: mouse | 95% C.I. = [0.35765, 0.75611] | into-L time points) | |
| 3A | zone/spd/zone encoding relation to transition | Chi Square | X2(4, n=1567) =137.3, p<0.0001 | n=1567 | neuron |
| 3B | DP relation to transition activation | Chi Square | X2(1, n=1565) =24.33, p<0.0001 | n=1565 | neuron |
| 4D | xCorr in zone transition, sepw | | | | |
| | GPe speed corr into L | | 95% C.I. of mean= [-0.09034, -0.02783] | n=9 | moouse |
| | GPe speed corr into D | | 95% C.I. of mean = [-0.1225, 0.02213] | n=9 | moouse |
| | GPe accel. corr into L | | 95% C.I. of mean= [-0.06978, 0.01473] | n=9 | moouse |

| | | | | | |
|---|---|---|---|---|---|
| | GPe accel. corr into D | | 95% C.I. of mean = [-0.09861, 0.05434] | n=9 | moouse |
| | GPi speed corr into L | | 95% C.I. of mean= [0.07538, 0.2694] | n=9 | moouse |
| | GPi speed corr into D | | 95% C.I. of mean = [-0.06794, 0.1414] | n=9 | moouse |
| | GPi accel. corr into L | | 95% C.I. of mean= [-0.04661, 0.01468] | n=9 | moouse |
| | GPi accel. corr into D | | 95% C.I. of mean = [-0.1172, 0.06086] | n=9 | moouse |
| | SNr speed corr into L | | 95% C.I. of mean= [0.05948, 0.2624] | n=8 | moouse |
| | SNr speed corr into D | | 95% C.I. of mean = [0.04388, 0.2848] | n=8 | moouse |
| | SNr accel. corr into L | | 95% C.I. of mean= [- | n=8 | moouse |

| | | | | | |
|---|---|---|---|---|---|
| | | | 0.2433, - 0.05349] | | |
| | SNr accel. corr into D | | 95% C.I. of mean = [- 0.1916, - 0.04448] | n=8 | moouse |
| 4M | Inter event interval between zones, SNr | paired ttest | p=0.0001 | n=8 | moouse |
| 4O | xCorr in zone transition, calb | | | | |
| | SNr speed corr into L | | 95% C.I. of mean= [- 0.02322, 0.08374] | n=8 | moouse |
| | SNr speed corr into D | | 95% C.I. of mean = [- 0.1154, 0.09804] | n=8 | moouse |
| | SNr accel. corr into L | | 95% C.I. of mean= [- 0.04114, 0.1098] | n=8 | moouse |
| | SNr accel. corr into D | | 95% C.I. of mean = [- 0.05105, 0.06158] | n=8 | moouse |
| 4R | Inter event interval between | paired ttest | p=0.0039 | n=7 | moouse |

| | | | | | |
|---|---|---|---|---|---|
| | zones, SNr, Calb | | | | |
| 5C | DS mCherry (subregion x MOR1) | 2-way RM ANOVA | F (2, 12) = 1.932, p=0.1874 | n=3 Ctrl; n=5 Gq | subregional average per mouse from 4-6 sections |
| | DS mCherry (subregion) | | F (1.516, 9.095)= 0.6642, p=0.4977 | n=3 subregions | |
| | DS mCherry (group: Ctrl, Gq) | | F (1, 6) = 0.01962, p=0.8932 | | |
| | DS mCherry (subject) | | F (6, 12) = 4.874, p=0.0096 | | |
| | DS MOR1 (subregion x MOR1) | 2-way RM ANOVA | F (2, 12) = 1.928, p=0.1879 | n=3 Ctrl; n=5 Gq | |
| | DS MOR1 (subregion) | | F (1.570, 9.419)= 6.124, p=0.0243 | n=3 subregions | |
| | DS MOR1 (group: Ctrl, Gq) | | F (1, 6) = 0.9410, p=0.3695 | | |
| | DS MOR1 (subject) | | F (6, 12) = 0.7792, p=0.6018 | | |

| | | | | | | |
|---|---|---|---|---|---|---|
| 5D | % time in dark (Ctrl vs Gq) | 2-tailed Mann-Whitney | p=0.9161 | n=17 Ctrl; n=20 Gq | mouse | Ctrl 72.52%±2.719; Gq 65.55%±5.556 |
| 5E | avg. spd cm/s (group x lighting) | within subjects | F (1, 35) = 2.147, p=0.1518 | n=17 Ctrl; 20 Gq | mouse | |
| | avg. spd cm/s (group: Ctrl, Gq) | 2-way RM ANOVA | F (1, 35) = 6.048, p=0.0190 | | | |
| | avg. spd cm/s (lighting: L, D) | | F (1, 35) = 47.34, p<0.0001 | | | |
| | avg. spd cm/s (subject) | | F (35, 35) = 3.229, p=0.0004 | | | |
| | post hoc (Ctrl L-vs-D) | Wilcoxon signed rank | p=0.0001 | n=17 Ctrl-L; n=17 Ctrl-D | mouse | Ctrl-L 6.152±0.2654; Ctrl-D 3.684±0.4485 |
| | post hoc (Gq L-vs-D) | Wilcoxon signed rank | p=0.0037 | n=20 Gq-L; n=20 Gq-D | mouse | Gq-L 4.412±0.5573; Gq-D 2.811±0.348 |
| | post hoc (Ctrl-L vs Gq-L) | 2-tailed Mann-Whitney | p=0.0044 | n=17 Ctrl; n=20 Gq | mouse | see above |
| | post hoc (Ctrl-D vs Gq-D) | 2-tailed Mann-Whitney | p=0.0417 | n=17 Ctrl; n=20 Gq | mouse | see above |
| 5F | maximum spd cm/s (Ctrl vs Gq) | 2-tailed Mann-Whitney | p=0.01 | n=17 Ctrl; n=20 Gq | mouse | Ctrl 22.18±0.8585; Gq 17.7±1.183 |

| | | | | | | |
|---|---|---|---|---|---|---|
| 5G | %time ≤2cm/s (Ctrl L-vs-D) | paired ttest | p<0.0001 | n=17 Ctrl-L; n=17 Ctrl-D; | mouse | Ctrl-L 32.18±2.123; Ctrl-D 50.18±3.561 |
| | %time ≤2cm/s (Gq L-vs-D) | paired ttest | p=0.0056 | n=20 Gq-L; n=20 Gq-D | | Gq-L 30.43±3.421; Gq-D 53.16±4.111 |
| | %time ≤2cm/s (Ctrl vs Gq in D) | Mann-Whitney | p=0.0059 | | | |
| | %time ≤2cm/s (Ctrl vs Gq in L) | Mann-Whitney | p=0.0168 | | | |
| 5H-I | approach (cm/s) impact of time | GLME of spd uing Matlab | 95% C.I. = [1.9925, 2.554] | n=1560 observations | time points | |
| & | approach (cm/s) impact of Gq | fitglme'; fixed effects: | 95% C.I. = [-0.89389, 1.0309] | | | |
| S 4 E,I | approach (cm/s) impact of Gi | | 95% C.I. = [-1.473, 1.0196] | | | |
| | approach (cm/s) impact of BIL | intercept, time, group, BIL; | 95% C.I. = [1.0773, 0.59214] | (17 Ctrl, 20 Gq, 15 Gi mice | | |
| | retreat (cm/s) impact of time | random effect: mouse | 95% C.I. = [-1.9511, -1.4538] | at 15 into-D and 15 into-L | | |
| | retreat (cm/s) impact of Gq | | 95% C.I. = [-1.507, 0.56654] | time points) | | |
| | retreat (cm/s) impact of Gi | | 95% C.I. = [- | | | |

| | | | | | | |
|---|---|---|---|---|---|---|
| | | | 1.3062, 0.92041] | | | |
| | retreat (cm/s) impact of BIL | | 95% C.I. = [1.0562, 1.486] | | | |
| | Gq approach (cm/s) impact of BIL | GLME of spd uing Matlab | 95% C.I. = [-0.13168, -1.1211] | n=600 observations | time points | |
| | Gq retreat (cm/s) impact of BIL | fitglme'; fixed effects: | 95% C.I. = [0.53604, 1.3761] | (20 mice @15 pnts app/ret) | | |
| | Gi approach (cm/s) impact of BIL | intercept, time, BIL; | 95% C.I. = [2.2463, 1.5553] | n=450 observations | time points | |
| | Gi retreat (cm/s) impact of BIL | random effect: mouse | 95% C.I. = [1.3261, 1.9938] | (15 mice @15 pnts app/ret) | | |
| | Ctrl approach (cm/s) impact of BIL | | 95% C.I. = [1.9076, 1.3185] | n=510 observations | time points | |
| | Ctrl retreat (cm/s) impact of BIL | | 95% C.I. = [0.9823, 1.615] | (17 mice @15 pnts app/ret) | | |
| 6C | % time in dark (Ctrl vs ChR) | 2-tailed Mann-Whitney | p=0.3941 | n=16 Ctrl; n=14 ChR | mouse | Ctrl 67.08±2.201; ChR 67.08±2.30 |
| 6D | avg. spd cm/s (lighting x group) | 2-way RM ANOVA | F (1, 26) = 1.687, p=0.2054 | n=30 (14 Ctrl, 16 ChR) | mouse | |

| | | | | | | |
|---|---|---|---|---|---|---|
| | avg. spd cm/s (group: Ctrl, ChR) | | F (1, 26) = 0.2461, p=0.6240 | | | |
| | avg. spd cm/s (lighting: L, D) | | F (1, 30) = 33.19, p<0.0001 | | | |
| | post hoc (Ctrl L-vs-D) | 2-tailed paired t-test | p<0.0001 | n=14 Ctrl-L; n=14 Ctrl-D | mouse | Ctrl-L 6.76±0.1801; Ctrl-D 4.76±0.2500 |
| | post hoc (ChR L-vs-D) | 2-tailed paired t-test | p=0.0011 | n=16 ChR-L; n=16 ChR-D | mouse | ChR-L 6.56±0.3170; ChR-D 5.195±0.2974 |
| | post hoc (Ctrl-L vs ChR-L) | 2-tailed Mann-Whitney | p=0.5979 | n=14 Ctrl-L; n=16 ChR-L | mouse | see above |
| | post hoc (Ctrl-D vs ChR-D) | 2-tailed Mann-Whitney | p=0.2803 | n=14 Ctrl-D n=16 ChR-D | mouse | see above |
| 6E | max spd cm/s (Ctrl vs ChR) | 2-tailed Mann-Whitney | p=0.4225 | n=14 Ctrl; n=16 ChR | mouse | Ctrl 23.308±0.692; ChR 24.625±1.341 |
| 6F | %time ≤2cm/s (Ctrl L-vs-D) | paired ttest | p=0.0002 | n=13 Ctrl; n=16 ChR | mouse | Ctrl-L 34.09±1.379; Ctrl-D 46.33±2.223 |
| | %time ≤2cm/s (ChR L-vs-D) | paired ttest | p=0.0005 | | | ChR-L 34.64±2.008; ChR-D 43.85±2.084 |
| | %time ≤2cm/s (Ctrl vs ChR in D) | unpaired ttest | p=0.4243 | | | |
| | %time ≤2cm/s | unpaired ttest | p=0.8305 | | | |

| | | | | | | |
|---|---|---|---|---|---|---|
| | (Ctrl vs ChR in L) | | | | | |
| 6M | % time in dark (Ctrl vs ChR) | 2-tailed Mann-Whitney | p=0.30629 | n=13 Ctrl; n=13 ChR | mouse | Ctrl 69.89±1.082; ChR 76.51±3.22 |
| 6N | avg. spd cm/s (lighting x group) | 2-way RM ANOVA | F (1, 24) = 0.0843, p=0.7740 | n=26 (13 Ctrl, 13 ChR) | mouse | |
| | avg. spd cm/s (group: Ctrl, ChR) | | F (1, 24) = 0.3329, p=0.5694 | | | |
| | avg. spd cm/s (lighting: L, D) | | F (1, 24) = 15.05, p=0.0007 | | | |
| | post hoc (Ctrl L-vs-D) | 2-tailed paired t-test | p<0.0001 | n=13 Ctrl-L; n=13 Ctrl-D | mouse | Ctrl-L 7.55±0.5792; Ctrl-D 5.328±0.5653 |
| | post hoc (ChR L-vs-D) | 2-tailed paired t-test | p=0.0012 | n=13 ChR-L; n=13 ChR-D | mouse | ChR-L 8.102±0.6756; ChR-D 5.510±0.6864 |
| | post hoc (Ctrl-L vs ChR-L) | 2-tailed Mann-Whitney | p=0.5403 | n=13 Ctrl-L; n=13 ChR-L | mouse | see above |
| | post hoc (Ctrl-D vs ChR-D) | 2-tailed Mann-Whitney | p=0.6139 | n=13 Ctrl-D n=13 ChR-D | mouse | see above |
| 6O | max spd cm/s (Ctrl vs ChR) | 2-tailed Mann-Whitney | p=0.1323 | n=13 Ctrl; n=13 ChR | mouse | Ctrl 24.154±0.986; ChR 27.385±1.824 |
| 6P | %time ≤2cm/s | paired ttest | p<0.0001 | n=13 Ctrl; n=13 ChR | mouse | Ctrl-L 30.29±2.633; |

| | | | | | | |
|---|---|---|---|---|---|---|
| | (Ctrl L-vs-D) | | | | | Ctrl-D 44.28±3.198 |
| | %time ≤2cm/s (ChR L-vs-D) | paired ttest | p=0.2460 | | | ChR-L 26.74±2.831; ChR-D 32.68±5.219 |
| | %time ≤2cm/s (Ctrl vs ChR in D) | unpaired ttest | p=0.0702 | | | |
| | %time ≤2cm/s (Ctrl vs ChR in L) | unpaired ttest | p=0.3684 | | | |
| S 2D | % time in dark (minnieScope mice) | summary statistics | | n=10 | mouse | 70.63%±4.613 |
| S 2E | average spd cm/s (minnieScope mice) | Wilcoxon signed rank | p=0.0098 | n=10 | mouse | L 5.46±0.4362; D 3.429±0.4308 |
| S 2F | max spd cm/s (minnieScope mice) | summary statistics | | n=10 | mouse | 15.9±0.9826 |
| S 2G | %time ≤2cm/s (L-vs-D) | Wilcoxon signed rank | p=0.0137 | n=10 | mouse | L 26.05±4.706; D 49.22±5.575 |
| S 2H-I | approach (cm/s) impact of time | GLME of spd uing Matlab | 95% C.I. = [0.0871, 0.6849] | n=300 observations | time points | |
| | approach (cm/s) impact of BIL | fitglme'; fixed effects: | 95% C.I. = [0.2506, -0.26603] | (10 mice with minniescope | | |
| | retreat (cm/s) impact of time | intercept, time, BIL; | 95% C.I. = [-1.9373, -1.2134] | at 15 into-D and 15 into-L | | |

| | | | | | | |
|---|---|---|---|---|---|---|
| | retreat (cm/s) impact of BIL | random effect: mouse | 95% C.I. = [1.576, 0.95053] | time points) | | |
| S 2K | zone encoding relation to speed encoding | Chi Square | X2(2, n=1565) =38.38 ,p<0.0001 | n=1565 | neuron | |
| S 3F | Inter event interval between zones, GPi | paired ttest | p=0.0018 | n=8 | moouse | L 6.482±0.5956; D 3.359±0.2066 |
| S 3K | Inter event interval between zones, Gpi | paired ttest | p=0.0035 | n=9 | moouse | L 7.969±0.9356; D 3.809±0.3087 |
| S 3K | Inter event interval between zones, Str,GRIN | paired ttest | p=0.0373 | n=8 | moouse | L 22.08±4.903; D 12.21±1.786 |
| S 4A | % time in dark (Ctrl vs Gi) | 2-tailed Mann-Whitney | p=0.8232 | n=17 Ctrl; n=15 Gi | mouse | Ctrl 72.52%±2.719; Gi 72.73%±2.719 |
| S 4B | avg. spd cm/s (group x lighting) | within subjects | F (1, 30) = 0.5551, p=0.4620 | n= (17 Ctrl, 15 Gi) | mouse | |
| | avg. spd cm/s (group: Ctrl, Gi) | 2-way RM ANOVA | F (1, 30) = 0.03006, p=0.8635 | | | |
| | avg. spd cm/s (lighting: L, D) | | F (1, 30) = 134.3, p<0.0001 | | | |
| | avg. spd cm/s (subject) | | F (30, 30) = 7.414, | | | |

| | | | | | | |
|---|---|---|---|---|---|---|
| | | | p<0.0001 | | | |
| | post hoc (Ctrl L-vs-D) | Wilcoxon signed rank | p=0.0001 | n=17 Ctrl-L; n=17 Ctrl-D | mouse | Ctrl-L 6.152±0.2654; Ctrl-D 3.684±0.4485 |
| | post hoc (Gi L-vs-D) | Wilcoxon signed rank | p<0.0001 | n=15 Gi-L; n=15 Gi-D | mouse | Gi-L 6.429±0.5924; Gi-D 3.622±0.5385 |
| S 4C | maximum spd cm/s (Ctrl vs Gi) | 2-tailed Mann-Whitney | p=0.715 | n=17 Ctrl; n=15 Gi | mouse | Ctrl 22.18±0.8585; Gq 21.6±1.245 |
| S 4D | %time ≤2cm/s (Ctrl L-vs-D) | paired ttest | p<0.0001 | n=17 Ctrl-L; n=17 Ctrl-D; | mouse | Ctrl L 32.18±2.123; Ctrl D 50.18±3.561 |
| | %time ≤2cm/s (Gi L-vs-D) | paired ttest | p<0.0001 | n=15 Gi-L; n=15 Gi-D | | Gi L 30.43±3.421; Gi D 53.16±4.111 |
| | %time ≤2cm/s (Ctrl vs Gi in D) | unpaired ttest | p=0.6601 | | | |
| | %time ≤2cm/s (Ctrl vs Gi in L) | unpaired ttest | p=0.5817 | | | |
| S 4F | avg. spd cm/s (group x lighting) | 2-way RM ANOVA | F (1, 27) = 0.06781, p=0.7965 | n=29 (16 Ctrl, 13 Gi) | mouse | Ctrl-LL 4.08±0.2428; Ctrl-DD 4.398±0.2145 |
| | avg. spd cm/s (group: Ctrl, Gi) | | F (1, 27) = 2.994, p=0.095 | | | Gi-LL 5.62±1.019; Gi-DD 6.037±0.9514 |
| | avg. spd cm/s (lighting: LL, DD) | | F (1, 27) = 3.651, p=0.0667 | | | |

| | | | | | | |
|---|---|---|---|---|---|---|
| | avg. spd cm/s (subject) | | F (27, 27) = 22.81, p<0.0001 | | | |
| S 4G | max spd cm/s (group x lighting) | 2-way RM ANOVA | F (1, 27) = 0.009815, p=0.9218 | n=29(16 Ctrl, 13 Gi) | mouse | Ctrl-LL 17.13±1.08; Ctrl-DD 17.25±1.112 |
| | max spd cm/s (group: Ctrl, Gi) | | F (1, 27) = 1.402, p=0.2467 | | | Gi-LL 19.92±1.806; Gi-DD 20.38±1.838 |
| | max spd cm/s (lighting: LL, DD) | | F (1, 27) = 0.05506, p=0.8163 | | | |
| | max spd cm/s (subject) | | F (27, 27) = 11.05, p<0.0001 | | | |
| S 4J | avg. spd cm/s (group: Ctrl, Gq) | 2-way RM ANOVA | F (1, 33) = 1.343, p=0.2549 | n=35(16 Ctrl, 19 Gq) | mouse | Ctrl-LL 4.08±0.2428; Ctrl-DD 4.398±0.2145 |
| | avg. spd cm/s (lighting: LL, DD) | | F (1, 33) = 2.676, p=0.1114 | | | Gq-LL 3.574±0.2999; Gq-DD 3.417±0.2953 |
| | avg. spd cm/s (group x lighting) | | F (1, 33) = 0.1536, p=0.6976 | | | |
| | avg. spd cm/s (subject) | | F (33, 33) = 4.919, p<0.0001 | | | |

| | | | | | | |
|---|---|---|---|---|---|---|
| S 4K | max spd cm/s (group: Ctrl, Gq) | 2-way RM ANOVA | F (1, 33) = 0.0007523, p=0.9783 | n=35(16 Ctrl, 19 Gq) | mouse | Ctrl-LL 17.13±1.08; Ctrl-DD 17.25±1.112 |
| | max spd cm/s (lighting: LL, DD) | | F (1, 33) = 2.683, p=0.1109 | | | Gq-LL 13.95±1.598; Gq-DD 15.47±1.294 |
| | max spd cm/s (group x lighting) | | F (1, 33) = 0.01158, p=0.9150 | | | |
| | max spd cm/s (subject) | | F (33, 33) = 4.179, p<0.0001 | | | |
